# Persistent Activity during Working Memory from Front to Back

**DOI:** 10.1101/2021.04.24.441274

**Authors:** Clayton E. Curtis, Thomas C. Sprague

## Abstract

Working memory (WM) extends the duration over which information is available for processing. Given its importance in supporting a wide-array of high level cognitive abilities, uncovering the neural mechanisms that underlie WM has been a primary goal of neuroscience research over the past century. Here, we critically review what we consider the two major ‘arcs’ of inquiry, with a specific focus on findings that were theoretically transformative. For the first arc, we briefly review classic studies that led to the canonical WM theory that cast the prefrontal cortex (PFC) as a central player utilizing persistent activity of neurons as a mechanism for memory storage. We then consider recent challenges to the theory regarding the role of persistent neural activity. The second arc, which evolved over the last decade, stemmed from sophisticated computational neuroimaging approaches enabling researchers to decode the contents of WM from the patterns of neural activity in many parts of the brain including early visual cortex. We summarize key findings from these studies, their implications for WM theory, and finally the challenges these findings pose. A comprehensive theory of WM will require a unification of these two ‘arcs’ of research.

## Introduction

The ability to store information for brief periods of time, so-called working memory (WM), is a building block for most of our higher cognitive functions, and its dysfunction is at the heart of a variety of psychiatric and neurologic symptoms. In the history of study into the neural mechanisms that support WM, an imperative goal of neuroscience, we would argue that there have been two main arcs. One began almost 50 years ago when Joaquin Fuster first reported that spiking measured from neurons in the macaque prefrontal cortex persisted during a WM delay (Fuster and Alexander, 1971). Following this seminal publication, many researchers have measured persistent activity with the goal of understanding how WM representations are stored by neural activity (Curtis and D’Esposito, 2003). The vast majority of the work has been focused on the prefrontal cortex. The other arc began more recently, over the last decade, but has already made a tremendous impact on WM theory. Utilizing sophisticated computational neuroimaging approaches (e.g., machine learning, encoding models, etc.), researchers demonstrated that one can decode the contents of WM from the patterns of neural activity in early visual cortex (e.g.,(Harrison and Tong, 2009; Serences et al., 2009). This was surprising because at the time no existing data, and surely no WM theory, suggested that sensory cortices played a role in WM storage. The so-called *sensory recruitment theory of WM* emerged from the ever-growing body of research suggesting a potential role for early visual cortex in visual WM.

## Neural activity persists in the prefrontal cortex

Following a century of studies investigating the effects of experimental lesions of the non-human primate cortex, researchers honed in on the principal sulcus in lateral PFC (from here on we will simply refer to this region as PFC) as a critical structure supporting WM functions (for a review see (Curtis and D’Esposito, 2004). By 1971, an American lab (Fuster and Alexander, 1971) and a Japanese lab (Kubota and Niki, 1971) began recording extracellular neurophysiological signals from the PFC while macaques performed WM experiments. They reported that some neurons in the PFC tended to maintain an elevated rate of spiking, relative to pre-trial baseline firing rates, during WM retention intervals. Adapting an oculomotor version of the delayed response task, along with other experimental refinements, allowed Funahashi, Bruce, and Goldman-Rakic (Funahashi et al., 1989) to clarify several features of the persistent activity. First, they demonstrated that persistent activity in PFC neurons was memory stimulus selective in that, for a given neuron, it was typically restricted to one or two of the target positions in the contralateral hemifield (Figure 1A). This meshed well with a later report that experimental lesions of the PFC tended to impact memory for targets in the contralesional hemifield (Funahashi et al., 1993a). Second, they demonstrated that activity persisted for the duration of the memory delay (3 or 6 seconds) consistent with a mechanism that bridged the time between the past sensory event and the contingent behavior. Third, they demonstrated that the amplitude of persistent activity was reduced prior to memory errors. Because these features align with our notions of memory so closely, persistent activity was embraced as the neural basis of working memory. It is no wonder, then, that the discovery of persistent activity is considered the most important scientific observation with regard to the neural mechanisms of working memory. This now classic finding has been replicated numerous times and has had a tremendous impact on WM theory and how we study WM experimentally (Riley and Constantinidis, 2015).

**Figure 1:**
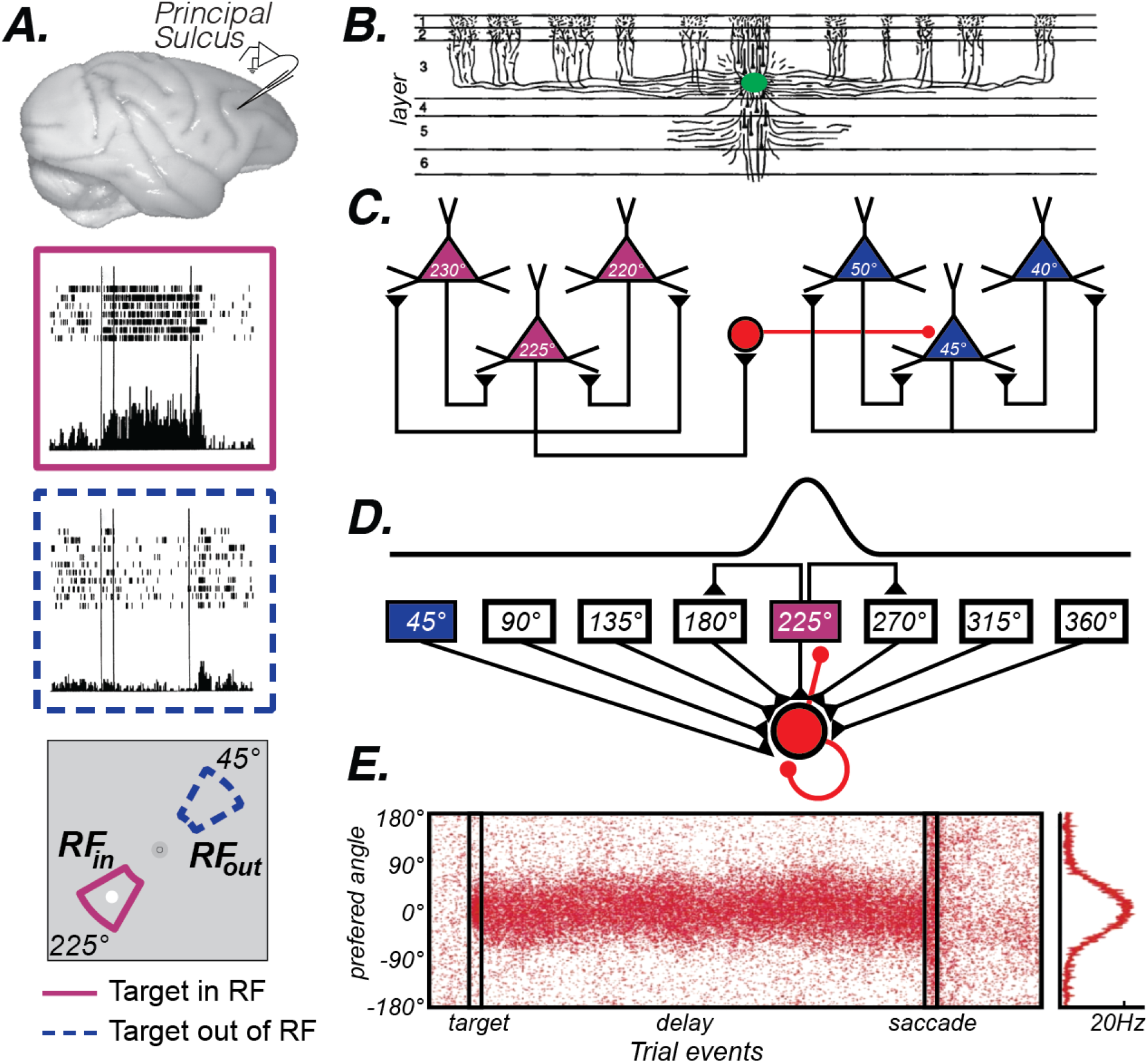
The canonical PFC microcircuit model of WM. A. Neural activity recorded from the principal sulcus in the macaque dorsolateral PFC. Activity persists during the delay period of memory-guided saccade tasks. The two insets depict a PFC neuron’s response when the memory target appears in and outside of it’s receptive or “memory” field. Adapted from data from Fig. 3 of (Funahashi et al., 1989). B. Tracers injected into deep Layer III of the macaque PFC (green blob) revealed the extensive lateral connections of pyramidal cells (Levitt et al., 1993). C. Goldman-Rakic (1995) hypothesized that these connections reflected similarly tuned pyramidal neurons, whose reciprocal excitatory and inhibitory connections enabled persistent activity. For instance, when remembering a target at 225° polar angle, recurrent excitation among similarly turned pyramidal neurons (purple triangles) maintains the location in WM through persistent activity. Inhibitory interneurons (red circle) suppress activity in neurons tuned to far away locations (blue triangles). Adapted from (Wang et al., 2013). D. This hypothesis was formulated into a computational theory in which both recurrent excitatory and inhibitory interactions were modeled (Wang, 2001) E. This model produces location-specific persistent activity similar to that observed in recordings of neurons in macaque PFC. Each red dot is a synthetic “spike” in a population of neurons with different location preferences, where the target is aligned at 0°. Location is encoded in the population response and decoding involves a read-out of the peak at any given time point. Adapted from (Compte et al., 2000).

Following these pioneering studies, the experimental techniques matured and over the next 30 years our knowledge about the relationships between persistent activity and WM accumulated. For example, persistent activity in PFC neurons is not limited to spatial WM. PFC neurons that show preferences for both simple (e.g., color) and complex (e.g., face) objects persist while monkeys maintain these objects in WM (Quintana et al., 1988; Miller et al., 1996; Ó Scalaidhe et al., 1999; Fuster et al., 2000). Assuming that stimulus selective persistent activity is the mechanism by which WM representations are stored, PFC neurons appear to store any type of stimulus feature, including the frequency of tactile flutter (Romo et al., 1999), the direction of dot motion (Zaksas and Pasternak, 2006; Mendoza-Halliday et al., 2014), sound location (Fuster et al., 2000; Kikuchi-Yorioka and Sawaguchi, 2000), and audiovisual macaque vocalizations (Hwang and Romanski, 2015). Moreover, they encode memory-guided prospective motor plans (Funahashi et al., 1993b; Takeda and Funahashi, 2002; Markowitz et al., 2015) and the prospective sensory features of a delayed paired associate (Rainer et al., 1999; Fuster et al., 2000). Finally, persistent activity appears to even encode complex task rules and contexts (Asaad et al., 2000; Wallis et al., 2001) and selective conjunctions of objects and locations (Rao et al., 1997; Rainer et al., 1998) that cannot be explained by simpler stimulus or location specific representations.

## Canonical PFC microcircuit model of WM

Once the link between persistent activity in the PFC and WM was firmly established, many focused on what were the properties of neurons and circuits in the PFC that give rise to memory selective persistent activity. Pyramidal neurons in layer III of the PFC make horizontal connections with clusters of other pyramidal neurons in regular intervals (Levitt et al., 1993; Lund et al., 1993; Kritzer and Goldman-Rakic, 1995) (Figure 1B). V1 neurons have a similar patchy horizontal connectivity (Gilbert and Wiesel, 1983) and connected neurons are more likely to have similar orientation tuning (Gilbert and Wiesel, 1989). By logic of induction, from these observations Goldman-Rakic theorized that similarly tuned (i.e., for location) pyramidal neurons in layer III are the source of glutamatergic excitatory recurrent connections that give rise to persistent activity (Goldman-Rakic, 1995) (Figure 1C). Indeed, the persistent activity of PFC neurons with similar visuospatial tuning are correlated (Constantinidis et al., 2001). These excitatory dynamics are thought to be balanced by closely synchronized fast spiking inhibitory interneurons (Constantinidis and Goldman-Rakic, 2002), whose lateral inhibition is theorized to additionally help sculpt the spatial tuning of PFC pyramidal neurons (Rao et al., 2000). Goldman-Rakic’s theory was formalized into a computational model that specified how excitatory recurrent activity, balanced and tuned by inhibition, could give rise to memory-specific persistent activity within a PFC microcircuit (Compte et al., 2000; Wang, 2001) (Figure 1D). This theoretical model highlighted the importance of the slow kinetics of NMDA receptors, compared to the faster kinetics of AMPA receptors (Wang, 1999). Empirical evidence has generally supported many aspects of the PFC microcircuit model of WM. Persistent activity depends on glutamatergic synapses on long, thin spines connecting PFC neurons in layer III (Wang et al., 2011), and these excitatory currents depend on the slow kinetics of NMDA receptors to support persistent activity (Wang et al., 2013). Moreover, the model hypothesizes that small random drifts in the bumps of activity cause the seemingly random inaccuracies in memory (Standage and Paré, 2018). Evidence for this hypothesis exists, as clockwise or counterclockwise biases in population estimates of delay activity in macaque PFC neurons predict small angular errors in memory (Wimmer et al., 2014).

There are also anatomical properties that suggest advantages that PFC may have in its capacity for WM storage. These slow NMDA receptors are densely expressed in PFC, especially when compared to V1 (Wang et al., 2008). Pyramidal neurons in PFC, again compared to visual cortex, have larger and more complex dendritic branching with a greater number of spines (Oga et al., 2017), and have more extensive horizontal collaterals in Layers II and III (Kritzer and Goldman-Rakic, 1995). Together, these anatomical features may better equip PFC with an increased capacity to integrate inputs, including the excitatory connections theorized to form positive feedback loops to sustain WM representations (Goldman-Rakic, 1995).

## Translating the primate PFC model of human WM

The success and impact of any animal model of human cognition depends on how well it translates to the species it is meant to model. It is not surprising then, that when brain imaging methods became widely available, researchers immediately predicted that indirect measures of neural activity could be used to measure persistent activity during WM in the homologous part of the human PFC. It turned out not to be so easy. The first human brain imaging study of spatial WM *failed* to find that blood flow measured with Positron Emission Tomography (PET) localized to the dorsolateral PFC (Jonides et al., 1993). Then, the failure of several studies to find spatial WM-related delay period activity in a homologous part of human dorsolateral PFC became the norm rather than the exception (Smith et al., 1996; Courtney et al., 1998; Zarahn et al., 1999; Rowe et al., 2000). A subsequent functional magnetic resonance imaging (fMRI) study from Goldman-Rakic’s own lab only succeeded in evoking dorsolateral PFC activity after increasing the WM load to 5 items (Leung et al., 2002). At the time it was assumed that fMRI did not have enough sensitivity to reliably measure persistent activity associated with maintaining a single item in WM. However, as we describe next this is unlikely the case and suggests alternative explanations.

Measuring neural activity with fMRI while humans perform spatial WM tasks, including memory-guided saccade WM tasks like those used to initially study the macaque PFC (Funahashi et al., 1989), we find that fMRI is perfectly sensitive to WM representations of single items (Curtis et al., 2004; Curtis and D’Esposito, 2006; Schluppeck et al., 2006; Srimal and Curtis, 2008; Tark and Curtis, 2009; Jerde et al., 2012; Sprague et al., 2014; Saber et al., 2015; Rahmati et al., 2020; Hallenbeck et al., 2021). However, in none of the above cited studies did we find evidence that neural activity persists in the human dorsolateral PFC during simple spatial WM tasks. On the other hand, in each one of those studies we found evidence that activity persists, in a variety of meaningful ways, in the superior spur of the precentral sulcus (PCS) in the frontal cortex and/or in the posterior part of the intraparietal sulcus (IPS) (Figure 2A).

**Figure 2:**
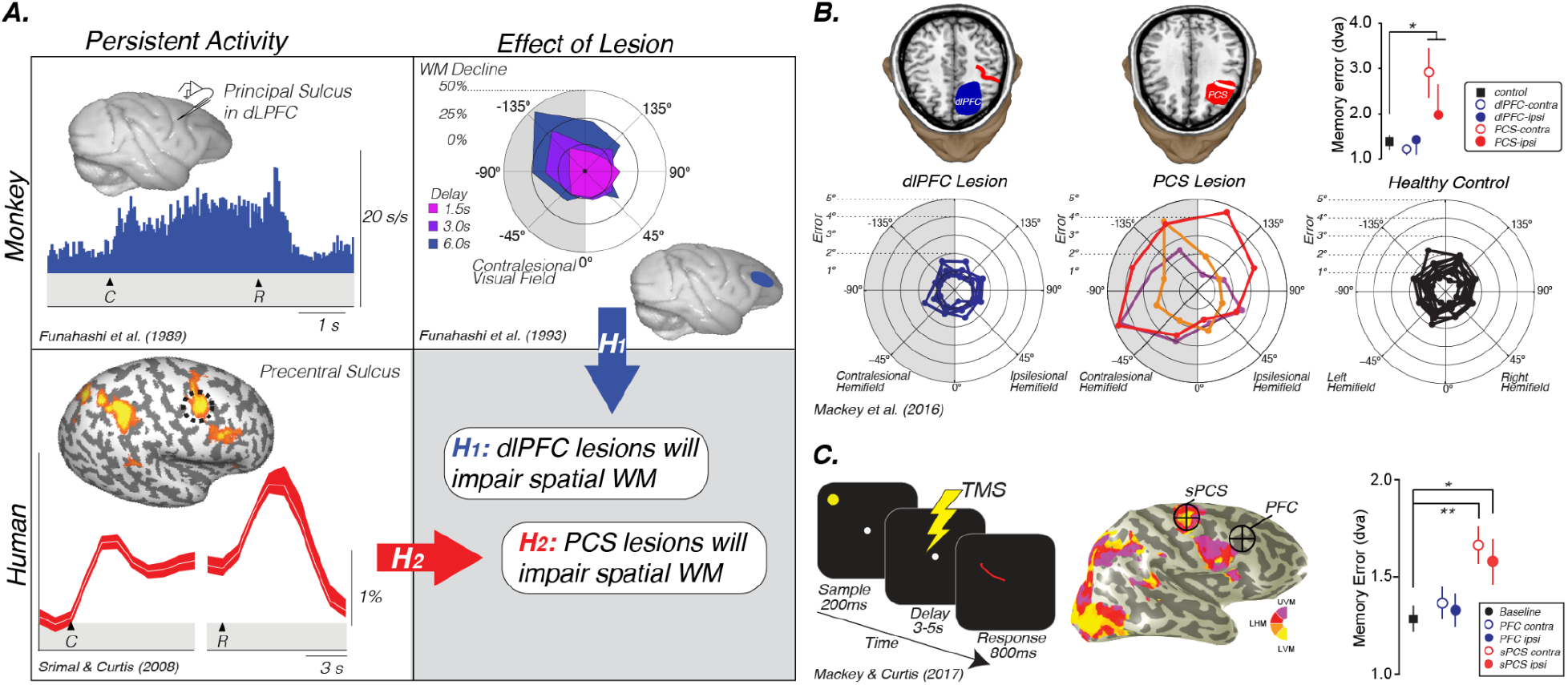
Translating the canonical model of WM to humans. A. Rationale and hypotheses of studies of lesion (Mackey et al., 2016b) and TMS perturbation (Mackey and Curtis, 2017) of human PFC. Neural activity persists in the monkey dlPFC during the retention interval of memory-guided saccade tasks (Funahashi et al., 1989). Lesions to the monkey dlPFC cause impaired memory-guided saccades, especially when made into the visual field contralateral to the lesion (Funahashi et al., 1993). Hypothesis 1: These monkey data predict that lesions to human dlPFC will impair spatial WM performance, including the accuracy of memory-guided saccades. However, human neuroimaging studies typically find persistent activity or multivoxel decoding of information restricted to the PCS, posterior to the likely homolog of the monkey principal sulcus in the dlPFC (Li et al.; Courtney et al., 1998; Srimal and Curtis, 2008; Jerde et al., 2012; Sprague et al., 2014; Hallenbeck et al., 2021). Hypothesis 2: These data predict that lesions to human PCS, not dlPFC, will impair WM performance. B. Human PCS lesions, but not dlPFC lesions, impact spatial WM (Mackey et al., 2016b). Plot in the upper right depicts the mean (SEM) of memory errors assessed by measuring the accuracy of memory-guided saccades when the memory targets were in the visual hemifield contralateral and ipsilateral to the lesion. The radial histograms show the spatial distribution of errors highlighting that the PCS lesions primarily impact memory-guided saccades to the contralesional hemifield, as in Funahashi et al., (1993). C. TMS applied during the middle of the delay period of a memory-guided saccade task to the retinotopically defined superior PCS, but not to dorsolateral PFC, induces errors in the accuracy of memory-guided saccades (Mackey and Curtis, 2017). Plot in the lower right depicts the mean (SEM) of memory errors assessed by measuring the accuracy of memory-guided saccades when the memory targets were in the visual hemifield contralateral and ipsilateral to the hemisphere in which TMS was applied.

On the face of it, these results conflict between the two species. In the monkey, neurons in dorsolateral PFC persist and lesions cause WM impairments. However, in humans neural activity only persists in the PCS, not in more anterior parts of the PFC in areas homologous to the macaque principal sulcus. We generated two hypotheses to explain these conflicting results based on the impact that lesions to the PFC and PCS had on WM (Figure 1A). If lesions to the dorsolateral PFC that spare the PCS cause WM impairments, like they do in monkeys, this would indicate that fMRI may not be sensitive enough to measure persistent activity in that part of the brain (hypothesis 1). On the other hand, if lesions to the PCS, rather than PFC, cause WM impairments, this would indicate that the human dorsolateral PFC is not necessary for WM like it is in the monkey (hypothesis 2). In support of hypothesis 2, the accuracy of memory-guided saccades was unimpacted by dorsolateral PFC resections as long as they spared the PCS (Figure 2B). PCS lesions increased the magnitude of memory errors largely when the target was in the contralesional hemifield (Mackey et al., 2016b, 2017; Mackey and Curtis, 2017)). In order to rule out other factors, like reorganization or compensation in the lesion patients, we repeated the study using transcranial magnetic stimulation (TMS) applied to the superior PCS and the intermediate frontal sulcus in the PFC during the memory delay in a healthy cohort of participants ((Mackey et al., 2016b, 2017; Mackey and Curtis, 2017)). The TMS results replicated the patient study; TMS to the sPCS, but not dorsolateral PFC, caused an increase in memory-guided saccade errors (Figure 2C). These results are consistent with previous studies that have investigated the impact of dorsolateral PFC and/or PCS damage on both spatial and non-spatial forms of WM (Ploner et al., 1999); (D’Esposito and Postle, 1999; Postle et al., 2003).

In understanding the discrepancy between these human and monkey studies, we need to consider a variety of possible explanations. First, a single item WM task may be too easy for humans relative to monkeys. Similar to the load argument discussed above (Leung et al., 2002), perhaps increasing the number of items increases the difficulty and thus recruits the human PFC. Nonetheless, the canonical WM theory does not specify that the dorsolateral PFC is only needed when the WM system is taxed with a challenging task. Moreover, other control processes such as reorganization and compression are necessary when one must maintain a number of items in WM (Rypma et al., 2002), especially when these approach or surpass capacity limits (Cowan, 2001). Second, the percentage of neurons in the macaque principal sulcus that show delay period activity is low (∼10%) relative to the percentage of neurons in the frontal eye field (FEF, ∼%50), located down in the anterior bank of the arcuate sulcus, and the lateral intraparietal area (LIP, ∼%50). Due to very large receptive fields, the tuning for location during the memory delay is coarse in the PFC relative to the FEF and LIP (Mohler et al., 1973; Blatt et al., 1990; Hamed et al., 2001). Plus, the horizontally connected clusters of pyramidal neurons in layer III of the PFC form stripes that are spaced 0.2 to 0.8 mm apart (Kritzer and Goldman-Rakic, 1995). Perhaps fMRI is insensitive because this spatial separation dilutes over voxels the signal from an already small percentage of poorly tuned neurons persisting in the PFC. Third, there are surely true differences between the two species that cannot be attributed to the methods with which neural activity is measured. If we were considering rodent models of WM (e.g., (Inagaki et al. 2019; Goard et al. 2016), we would be less bothered by possible mismatches in the exact brain areas, and would instead focus on the advantages of the animal model to learn about the precise neural mechanisms. One potential implication is that the mechanisms described in the microcircuit model of WM might be more applicable to cortical areas other than the human PFC. Indeed, lesions to the macaque FEF and LIP, as well as homologous areas in the human brain both impair WM performance (Dias and Segraves, 1999; Gaymard et al., 1999; Li et al., 1999; Ploner et al., 1999; Mackey et al., 2016a, 2016b; Mackey and Curtis, 2017).

## Neural activity persists beyond PFC

The dorsolateral PFC is not the only brain area housing neurons that persist during WM (Leavitt et al., 2017). Funahashi et al., (1989) also reported that neurons in the FEF showed spatially tuned persistent activity. As mentioned above, persistent activity is more common among FEF neurons than PFC, more robust, and more spatially selective (Goldberg and Bruce, 1985; Sommer and Wurtz, 2001; Merrikhi et al., 2017; Hart and Huk, 2020). Activity persists during WM tasks in several other frontal areas including the dorsal premotor cortex (PMD; (Rossi-Pool et al., 2017; Bastos et al., 2018), the supplementary eye fields (SEF; (Shichinohe et al., 2009; Fukushima et al., 2011), the anterior cingulate cortex (ACC; (Kamiński et al., 2017), and even the orbitofrontal cortex (OFC; (Ichihara-Takeda and Funahashi, 2007). Moreover, neurons in LIP and 7a also show spatially selective and robust persistent activity (Gnadt and Andersen, 1988; Barash et al., 1991; Constantinidis and Steinmetz, 1996; Chafee and Goldman-Rakic, 1998; Pesaran et al., 2002; Hart and Huk, 2020). In the temporal lobe, WM selective persistent activity has been reported in neurons in monkey inferotemporal (IT) cortex (Fuster and Jervey, 1981; Miyashita and Chang, 1988; Miller et al., 1993; Chelazzi et al., 1998) and even in hippocampus and nearby entorhinal/perirhinal cortex (Miller and Desimone, 1994; Suzuki et al., 1997; Wirth et al., 2003). Evidence also exists that persistent neuronal activity carries sensory information about object identity in V4 (Hayden and Gallant, 2013) and motion in area MT (Bisley et al., 2004). However, these results are controversial (Pasternak and Greenlee, 2005; Leavitt et al., 2017) as other studies have reported an absence of persistent activity among neurons in MT coding for the remembered motion direction (Mendoza-Halliday et al., 2014). The discrepancy may lie in the type of representational format an animal might be using to store the memory as opposed to the representation formed during perception. For instance, it is unlikely that memory for dot motion is a replay of hundreds of dots moving over time. Perhaps, that temporally evolving percept is compressed or recoded into something like a single directional vector that does not drive MT. Remarkably and surprising to many neuroscientists, even neurons in V1 show activity which persists during WM delays (Supèr et al., 2001; van Kerkoerle et al., 2017). Finally, neurons in subcortical areas like the superior colliculus (SC; (Shen et al., 2011; Dash et al., 2015; Sadeh et al., 2018) and mediodorsal thalamus (Funahashi, 2013) are spatially tuned and carry location information during WM delay periods. The point we are trying to make in this section is that the presumed mechanism that supports WM - memoranda-specific persistent activity - is not specific to the dorsolateral PFC. Rather it appears to be a mechanism used by many parts of the brain to encode enduring representations useful for memory-guided decisions.

In humans, measuring delay period activity with fMRI BOLD supports these reports in just how widely distributed persistent activity appears to be during WM. There have been a number of reviews recently of human neuroimaging studies of WM (e.g., (Sreenivasan et al., 2014; D’Esposito and Postle, 2015; Christophel et al., 2017; Sreenivasan and D’Esposito, 2019) and thus we will instead focus on instructive examples of persistent activity measured in humans with fMRI. In many of the monkey electrophysiological studies reviewed above, an important first step involved characterizing each neuron’s receptive field (RF) or its preferred stimulus feature. This then allowed researchers to compare memory responses between stimuli placed within and outside of each neuron’s RF (or compare between preferred and non-preferred stimuli). Utilizing the same logic, advances in population receptive field (pRF) mapping (Dumoulin and Wandell 2008; Mackey et al. 2017; Wandell and Winawer 2015) allow researchers to compare BOLD estimates of persistent activity between trials in which the memoranda fall within and outside of a voxel’s pRF. In Figure 3, the time courses of BOLD activity during a memory-guided saccade task are shown for ten visual field maps (Rahmati et al., 2020; Hallenbeck et al., 2021). Each visual field map contains either an upright or inverted representation of the contralateral visual field. The location and size of the pRF of each voxel in these maps can be estimated. Averaging BOLD signal over trials can be done in a principled way according to the match between voxels’ pRF and the locations of the memorized targets. Overall, activity persists during the delay period in almost all of these maps. Moreover, the amplitude of BOLD activity is generally greater among voxels with pRFs matching the target, compared to voxels with pRFs 180° away from the target (on the opposite side of fixation).

**Figure 3.**
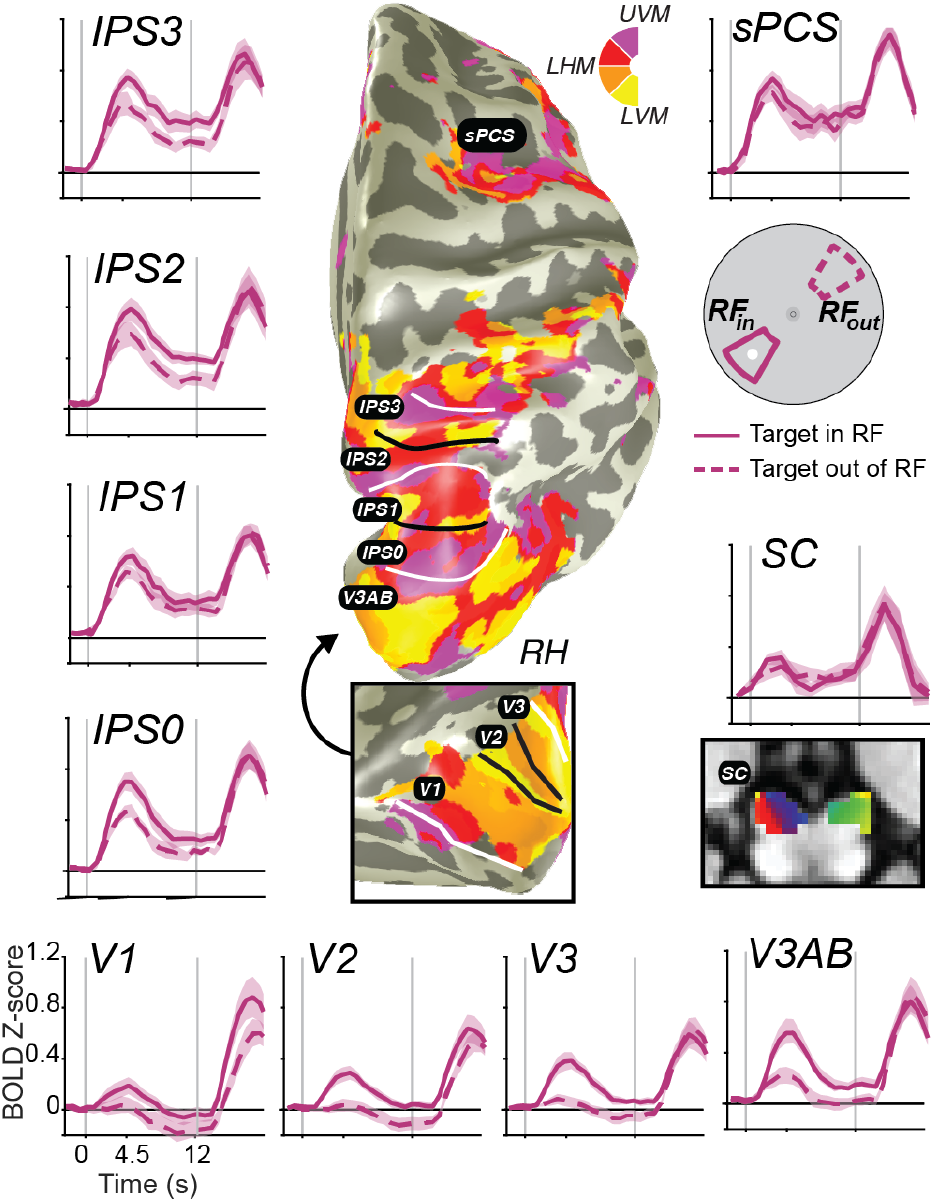
Stimulus-selective persistent activity across the human brain measured with fMRI. BOLD activity persists during the retention interval of memory-guided saccade tasks in many cortical and subcortical brain areas (Hallenbeck et al., 2021). Each region-of-interest was defined using modified population receptive field (pRF) mapping procedures (Mackey et al., 2017; Rahmati et al., 2020). Solid lines are the average BOLD signal from trials in which the memory target fell within the pRF of voxels. Dashed lines represent the averaged signal from trials in which the target was 180 degrees away from the target. Error bands are SEM. Gray vertical lines represent the onset of a brief visual target and the end of the retention interval. Notice how the general amplitude of persistent activity increases from early visual cortex to parietal cortex to frontal cortex, but the spatial selectivity generally decreases. Moreover, BOLD activity persists in retinotopically defined human superior colliculus (SC; (Mackey et al., 2017; Rahmati et al., 2020)).

Based on these data, two major gradients can be seen. One, the overall amplitude of persistent activity increases moving up the visual hierarchy from visual cortex to parietal cortex to frontal cortex. Two, the spatial selectivity (difference between when the target is in or out of the voxel’s pRF) generally decreases up that same hierarchy. Even within the parietal cortex, we can see both of these gradients from IPS0 to IPS3. Of special interest, many neuroimaging studies have failed to find persistent activity in V1 (or in early visual cortex for that matter; e.g., (Ester et al., 2009; Harrison and Tong, 2009; Offen et al., 2009; Serences et al., 2009; Riggall and Postle, 2012; Albers et al., 2013). These studies, however, typically averaged over all of V1, likely missing the more localized activity that persists. Note how the voxels in V1 with pRFs overlapping the small memory target showed a brief transient response time-locked to the target stimulus, but activation does not remain above the pre-trial baseline for the entire delay period. On the one hand, V1 does not meet the strict definition of persistent activity. On the other hand, if one considers the relationship between encoding and decoding from neural populations, then it does meet the definition of persistent activity. Namely, the WM representation is clearly encoded in the population as evident by the difference between the two time-courses. Moreover, the same decoder (e.g., max) applied to read-out the population response would recover the target location despite the average signal dipping back down to pre-trial levels. We reported the same pattern in V1 previously even on trials in which the location of the visual target was different from the location of the memory-guided saccade, using an antisaccade procedure (Saber et al., 2015), indicating that the response is memory related and not solely a residual BOLD response due to the visual transient. These advances allow for promising and more direct comparisons between monkey electrophysiology and human neuroimaging.

## Decoding WM contents from activation patterns: another form of persistent activity

As reviewed above, attempts to identify persistent activity in human PFC that is critical for behavioral performance have largely been unsuccessful. However, these studies typically leveraged mass univariate analysis approaches which primarily focus on the *similarities* in fMRI activation across particular remembered stimuli. That is - to isolate activation related to the maintenance of information over the delay period, trials corresponding to all possible WM contents are averaged. Such averaging necessarily masks important differences in activation associated with specific types of stimuli - for example, the particular location, orientation, or color held in WM.

Beginning at the turn of the century, human neuroimaging researchers began considering the possibility that *patterns* of brain activation measured with fMRI could discriminate between different stimulus or task conditions, rather than only considering elevated or suppressed *average* activation (Haxby et al., 2001; Norman et al., 2006). These methods primarily involve ‘decoding’ which of several stimuli was present using machine learning tools, such as support vector machines. When these methods were turned to the early visual system, they demonstrated a remarkable ability to decode which orientation was viewed based on visual cortex activation patterns (Haynes and Rees, 2005; Kamitani and Tong, 2005). This was a surprising feat - the anatomical organization of orientation selectivity in the early visual system was thought to be too fine for study with the relatively-coarse spatial resolution of fMRI (on the order of 2-3 mm per voxel). However, because the coarse sampling of the fine orientation columns is imperfect and uneven, the observed pattern of activation differed across stimulus orientations, enabling the decoding algorithm to detect these subtle differences and accurately decode which orientation was viewed (Boynton, 2005; Swisher et al., 2010). It should be noted that there exists considerable skepticism about the exact signals driving successful orientation decoding performance in these studies (Freeman et al., 2011, 2013; Alink et al., 2013; Carlson, 2014; Maloney, 2015; Pratte et al., 2016; Roth et al., 2018). Regardless of the source of the signals, it remains possible to recover distinctions in brain activation patterns associated with visual stimulus features.

Soon thereafter, these methods were applied to WM: Frank Tong (Harrison and Tong, 2009) and John Serences (Serences et al., 2009) each reported success applying similar decoding techniques to visual cortex fMRI activation patterns measured during the delay-period of WM tasks. In each case, the authors demonstrated that only remembered information could be decoded, and non-remembered information (e.g., a discarded feature (Serences et al., 2009), or a discarded stimulus (Harrison and Tong, 2009) was not maintained, demonstrating that these results cannot only be due to lingering sensory-evoked activation present in the slow hemodynamic signals measured with fMRI. This pair of studies offered convincing evidence for an important role of early sensory regions in supporting WM representations, especially when the features to be maintained are well-represented within those regions. This *sensory recruitment model* of WM posits that the previously-identified sustained delay-period activation observed in association cortex acts to coordinate stimulus-specific representations in sensory cortex (Curtis and D’Esposito 2003; Serences 2016; Postle 2006; D’Esposito and Postle 2015)

In the decade since, dozens of studies have applied similar methods to decode visual stimulus features such as orientation, motion direction, color, spatial position, and the identity of a spatial pattern from brain activation patterns measured from striate and extrastriate visual cortex (Christophel et al., 2017). Moreover, modified versions of these decoding methods, including cvMANOVA (Christophel et al. 2018; Allefeld and Haynes 2014), inverted encoding models (IEM; Fig. 4A; (Ester et al., 2013; Sprague et al., 2014)), and Bayesian decoding methods (Li et al.; van Bergen et al., 2015; van Bergen and Jehee, 2018, 2021; Brissenden et al., 2021) have increasingly improved the resolution and sensitivity of these methods to differences between conditions, and, ultimately, between individual trials. These new methods have revealed feature-selective representations broadly across visual, parietal, and frontal cortex (Li et al.; Christophel et al., 2012, 2018a, 2018b; Jerde et al., 2012; Christophel and Haynes, 2014; Ester et al., 2015; Yu and Shim, 2017; Rahmati et al., 2018), along with subcortical regions including the superior colliculus (Rahmati et al., 2020) and cerebellum (Brissenden et al., 2021). In human neuroimaging, evidence for stimulus-selective persistent activity abounds throughout the brain (Fig. 4B).

**Figure 4:**
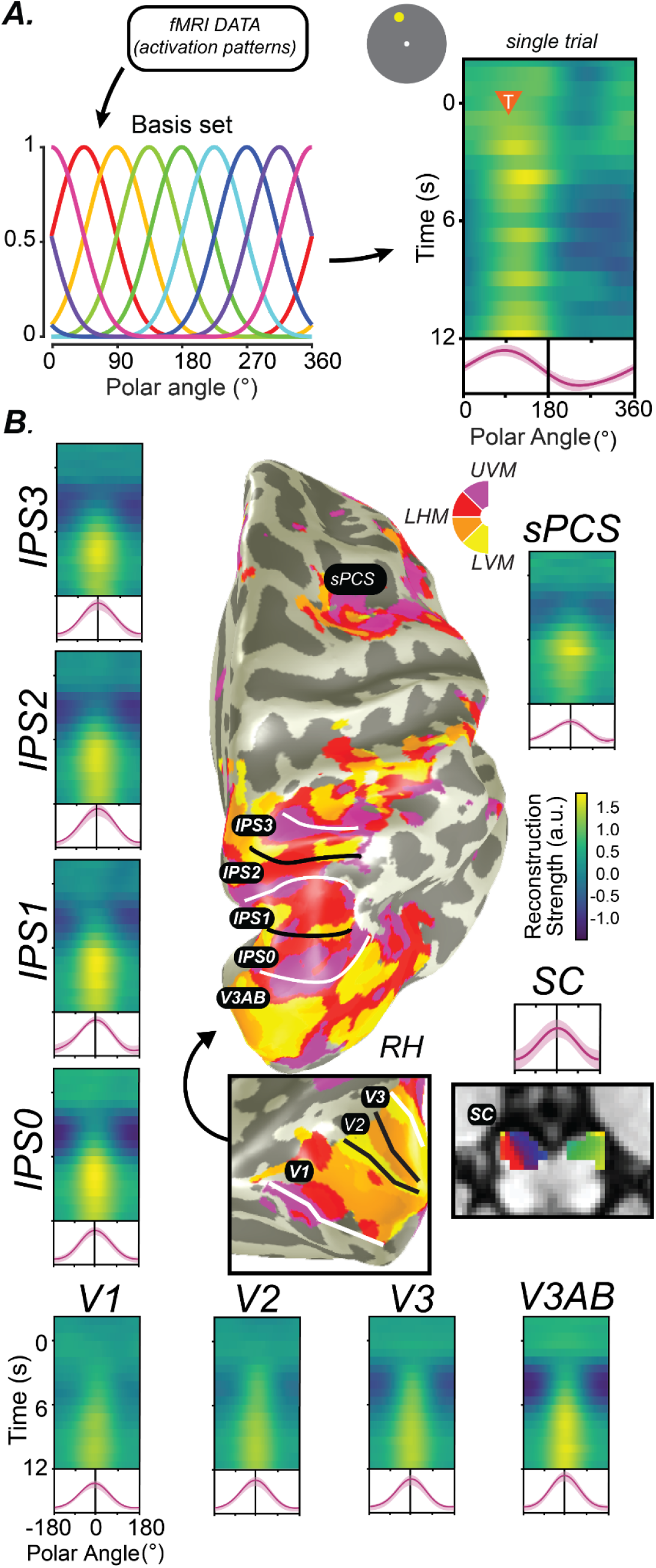
Decoding methods reveal stimulus-selective persistent activity across cortical and subcortical regions in the human brain. (A) Recent studies have employed ‘inverted encoding models’ (IEM), which model the activation of each voxel as a weighted combination of neural information channels (Brouwer and Heeger, 2009; Ester et al., 2013; Sprague et al., 2018). Applying this method results in reconstructed channel response profiles for each timepoint of each trial. When a measured activation pattern contains a representation of the remembered information, these reconstructed channel response profiles peak at the corresponding feature value. Right: single example trial illustrating a persistent representation of the remembered location on that trial based on activation patterns in V3AB (orange triangle indicates onset of delay period and remembered feature value). (B) When applied to activation patterns in retinotopic cortical regions and the superior colliculus while participants perform a memory-guided saccade task, representations of remembered positions are universally recovered. For each ROI, we show a timecourse of reconstructed representations (each row is a single timepoint; average of reconstructions for all experimental trials aligned to the remembered location), along with the average channel response profile over the final 1.5 s of the delay period (red line). Data adapted from Hallenbeck et al, (Hallenbeck et al., 2021) (A-B; all ROIs except SC) and Rahmati et al, (Rahmati et al., 2020) (B; SC). This Figure depicts the same data shown in Fig. 3 analyzed in a different way.

Interestingly, in many cases, when sustained delay-period activation is compared directly against stimulus-selective activation patterns, complementary results are found (Postle, 2015). As an example, Riggall & Postle (Riggall and Postle, 2012) compared univariate delay-period activation and decoded information content for regions responsive to visual stimuli and those with elevated responses during the delay period of a WM task. In the stimulus-responsive regions (which were primarily in extrastriate visual cortex), a decoding algorithm was able to successfully recover the direction of motion remembered by the participants, but these regions did not show elevated delay-period activation. Conversely, in delay period-responsive regions (which were primarily in the intraparietal sulcus dorsal frontal cortex), the authors could not decode the remembered stimulus value, but did observe sustained delay-period activation spanning the sample and the probe stimulus. In a subsequent study in which WM load was additionally manipulated, sustained delay-period activation in frontal and parietal regions was shown to increase as WM load increased from 1 to 3 items, while a similar change in average activation was not observed in sensory regions (Emrich et al., 2013). However, the authors could reliably decode the remembered stimuli from activation patterns in sensory regions, with accuracy decreasing as WM load increased. Once again, this suggests that regions showing elevated delay-period activity may not be those which represent the WM content itself, and that instead there may be a division of labor between frontal and parietal regions which help coordinate WM representations and sensory regions which encode stimulus values themselves.

However, when interpreting results from these decoding studies, it is critical to consider how these various algorithms operate to discriminate between remembered visual stimuli. The key feature of any decoding algorithm is that it identifies a reliable difference between activation patterns associated with different modeled stimulus values. While different approaches use different assumptions about the structure of these activation patterns and their noise covariance, this core feature remains. Accordingly, if a decoder can reliably pick up on differences between activation patterns within a region associated with different stimulus values, this necessarily means that some neurons (or, at least, signals resulting from neural activity) are more active than others in a reliable way. That is - the decoders aren’t magic - they’re just exploiting the structure of signals measured from neural tissue to optimally extract activation associated with different stimulus values. And, importantly, some stimulus values result in increased activation in some measured units, while other stimulus values result in increased activation in other measured units. As a trivial example, one could build a visual stimulus decoder based on a machine learning algorithm (e.g., support vector machines) to decode which side of the screen is stimulated by a large flickering checkerboard - a stimulus that is well-understood to evoke extremely strong and reliable fMRI signals in contralateral visual cortex. The decoder would perform extremely well - likely approaching 100% correct decoding performance. While in this case it wouldn’t be necessary to apply the decoding algorithm to show that primary visual cortex encodes the retinotopic location of a stimulus, because a simple fMRI contrast would reveal strong evidence for such a result, this example remains illustrative: the decoder would be basing its judgment on localized increases in activation within a subset of the population of voxels.

When such an analysis is applied to data acquired during a memory-guided saccade task analogous to that used in macaques, greater activation is measured in voxels with spatial RFs near the remembered location as compared to those voxels with spatial RFs farther away (Fig 3; (Saber et al., 2015; Hallenbeck et al., 2021). Moreover, this holds for features like orientation: recent studies which have instead attempted to ‘localize’ voxels preferring one or another orientation and directly compare activation between these subpopulations support this notion: voxels labeled with the orientation remembered on a trial show elevated activation compared with those labeled with the non-remembered orientation (Lawrence et al., 2018). These results track with those observed in the classical studies of macaque DLPFC which show elevated neural firing for neurons which prefer the remembered location as compared to those with more distal preferences (Funahashi et al., 1989).

In our view, *this is the very definition of persistent activity.* Thus, decoding studies which observe stimulus-selective activation patterns in different cortical and subcortical brain regions should be considered to provide support for stimulus-selective persistent activity.. That is - decoding of WM content and elevated delay-period activation may in many cases be considered two sides of the same coin (Fig. 3 vs Fig. 4B). Recent advances in decoding methods described above (IEM, cvMANOVA, and Bayesian generative models) have further extended the set of regions from which WM content can be decoded. Ester et al (2015) and Yu & Shim (2017) applied IEMs to decode orientation and color from several parietal and prefrontal regions, and Christophel et al (2018) applied a non-parametric decoder based on a multivariate ANOVA to decode remembered orientation from the same regions from a large sample of fMRI participants (n = 87). Thus, in many cases, where sustained delay-period activation is found in humans, successful decoding of WM content soon follows (as the capabilities of methods advance).

## Further challenges to the canonical PFC model of WM

So far we have described several findings that challenge the canonical WM model. In humans, simple WM does not depend on the dorsolateral PFC. Additionally, the persistent activity of neurons in PFC that sits at the heart of the canonical WM model is observed in many other brain regions, including early visual cortex. Together, these findings suggest that perhaps theories have tended to overemphasize the unique importance of PFC for WM. Additionally, further challenges have recently arisen to the very nature of what role persistent activity plays in WM.

### Persistent activity in PFC is an artifact of averaging

First, some have questioned whether persistent activity in PFC neurons is an artifact of averaging over trials (Shafi et al., 2007; Stokes and Spaak, 2016; Spaak et al., 2017). Similarly, the spiking activity of single PFC neurons might be best described as idiosyncratic bursts rather than persistent, and perhaps PFC activity is better characterized as ‘bubbles’ of oscillations in LFP (Lundqvist et al., 2016, 2018). However, even if one accepts this to be the case, the original theoretical model does not need to be adjusted. The canonical model put forth by Goldman-Rakic (1995) and its later formalization as a computational model (Compte et al., 2000) never specified that WM representations were stored by the persistent activity of single neurons. On the contrary, even the earliest versions of the model were inspired by the anatomy of layer III PFC neurons that were proposed to be clustered in pools of similarly tuned neurons with recurrent excitatory connections. Furthermore, the computational model clearly encodes WM representations through the overall activity of a population of neurons. Perhaps the fact that the evidence for the theory took the form of averaged recordings of single neurons may have confused the issue.

### Dynamic codes for WM content

Second, some have questioned the temporal stability of WM representations encoded by the delay period activity of PFC neurons (Parthasarathy et al., 2017, 2019; Spaak et al., 2017; Cavanagh et al., 2018; Wasmuht et al., 2018). Based on analyses comparing the activity of groups of neurons across timepoints within trials, these studies have concluded that the population activity of PFC neurons that code for WM representations dynamically changes over time. This could be a real challenge to the canonical model of WM because this mechanism is at odds with a stable fixed activity pattern linking neuronal turning preferences with features stored in WM. Specifically, if WM representations were primarily dynamic, a downstream area would have to know about and track the dynamics of each neuron’s encoding properties (its mnemonic tuning function as it unfolds over the trial) in order to read out the represented feature value from the population response at a given timepoint.

However, recent theoretical and empirical demonstrations have mitigated these concerns. Even when activity patterns are somewhat dynamic, such that the correlation between activity patterns is lower for points further separated in time than for points nearer in time, the population can be shown to have the same information content. Specifically, Murray et al (2017) demonstrated that dynamic activity patterns that are occasionally observed in PFC exist within a ‘stable subspace’ of the full population activity space, such that a downstream region could apply a fixed linear readout to accurately recover WM information throughout the delay period. Thus, at least in some cases, dynamic codes only appear this way on the surface (Murray et al., 2017; Parthasarathy et al., 2019).

While there certainly does exist ample evidence that dynamic responses at the single-unit level can be observed, and that they can in some cases support a stable population-level neural code, it is critical to note that these studies do not negate the existence nor importance of other stable coding mechanisms, some of which are observed in the same studies. For example, it has been shown that neurons with dynamic responses and those with stable responses coexist in PFC, and their response dynamics can be well-predicted by their intrinsic ‘time constant’ (their autocorrelation function measured from inter-trial intervals; (Wasmuht et al., 2018). That is - nearby neurons in the same brain region can either show evidence for dynamic coding or stable coding. In another study, macaques performed an oculomotor delayed response task with an intervening irrelevant distractor stimulus. Activity patterns measured from LPFC ‘morphed’ following the distractor, but patterns measured from the FEF of the same animals did not show evidence for such dynamic morphing (Parthasarathy et al., 2017). These results show that even when dynamic codes are observed, stable subspaces (consistent with a fixed readout rule) can account for a large amount of the response dynamics, and moreover, that stable coding is simultaneously observed in other neurons and/or brain regions.

### Mixed selectivity in PFC

Third, PFC neurons appear to have mixed selectivity as they can change their responsiveness to the same stimulus or behavioral response depending on subtle contextual changes within a task (Sigala et al., 2008; Machens et al., 2010; Mante et al., 2013; Rigotti et al., 2013). This could have several implications, including that the population response does not encode straightforward task variables, or that it encodes some latent variables that have yet to be discovered, or that it is dynamic over time at the timescale of the recording session. Nonetheless, there are some advantages to mixed selectivity. For example, the idea that the PFC can store any type of feature in WM implies that the entire manifold of encoding mechanisms housed in our sensory cortices might need to be duplicated just for short term storage, which seems highly inefficient at best. Mixed selectivity could vastly increase the encoding capacity of a given population (Rigotti et al. 2013). However, incorporating this concept into the canonical PFC model of WM would require altering the theory in ways that approach the way in which the hippocampus is thought to use mixed selectivity and sparse coding for long-term memory (Rolls and Treves, 1990; McClelland et al., 1995). Moreover, perhaps we have yet to discover the mechanisms by which the population response in PFC is demixed when it is readout by other brain areas (Machens, 2010).

### “Activity-silent” WM representations

Fourth, metabolically economical models propose that persistent spiking may induce fast-timescale synaptic changes that encode stimulus properties that can be later retrieved efficiently via stimulus-agnostic “pinging” of the network (Mongillo et al., 2008; Stokes et al., 2013; Rose et al., 2016; Wolff et al., 2017), instructive cues (Lewis-Peacock et al., 2012; Sprague et al., 2016; LaRocque et al., 2017; Lorenc et al., 2020), and/or spontaneous internal neural reactivation signals (Lundqvist et al., 2016, 2018). In the empirical reports, decoding performance reliably drops around chance levels at one point in the trial, but a subsequent visual stimulus (Wolff et al., 2015, 2017, 2020a, 2020b), TMS pulse (Rose et al., 2016), or task instruction (Lewis-Peacock et al., 2012; Sprague et al., 2016; LaRocque et al., 2017) results in a ‘reactivation’ of an otherwise ‘latent’ WM representation. This negative evidence, in the form of poor or at-chance decoding performance prior to reactivation, has been used to suggest that currently irrelevant information in WM is not maintained in an active state accessible to the neural signals fed into the decoding algorithm.

However, these studies additionally cannot rule out a key role for persistent activity in supporting WM behavior. When larger sample sizes and more sensitive analysis techniques are applied, there appears to be some positive evidence for representations of irrelevant WM information (Christophel et al., 2018b). While this one positive result does not invalidate the previous negative observations, it does suggest that it can be possible - with a sufficient sample size - to find evidence for WM representations that elude studies with smaller sample sizes. The studies which have ‘pinged’ human participants with irrelevant visual stimuli or TMS pulses can also not conclusively demonstrate that there existed *no information* prior to the reactivation stimulus. While the information may not have been accessible with EEG or MEG measurements, it may have existed as spontaneous oscillations in the electrophysiological recordings (LaRocque et al., 2013; Foster et al., 2016). A recent reanalysis of the data shown in Wolff et al (2017) suggests this latter possibility (Barbosa et al., 2021). Finally, modeling has shown that observations of increased information content in IEM-based stimulus reconstructions following a task cue (Sprague et al., 2016) are not diagnostic of a transition from a passive to an active code (Schneegans and Bays, 2017). While this study (Sprague et al., 2016) found an enhancement in the decodable information about remembered spatial position following an informative cue, it remains the case that weak information may have been present prior to the cue, but was inaccessible to the fMRI signal and/or decoding algorithm employed (e.g., Christophel et al., 2018b).

### Distractors impact WM representations in sensory regions

Cognitive theories about the nature of WM representations have long been informed by behavioral studies of the distracting effects of material presented during WM retention intervals (Baddeley, 1986). Intervening information is more disruptive when its features match the contents of WM. For instance, intervening phonological but not visual information impairs one’s ability to maintain visually presented strings of letters, suggesting an important role of articulatory processes for items that are verbalizable (Logie et al., 1990). Similarly, intervening visuospatial processing and oculomotion selectively impacts spatial WM (Postle et al., 2006). In general, many conclusions about the formats of WM representations depend on the logic that the effectiveness of distraction depends on how well the representational formats of the distractor and memoranda are matched. Neuroscientific studies have also relied on a similar logic, assuming that the competition between the neural representations of the memoranda and the intervening distractor disrupts memory. As reviewed above, early electrophysiology and neuroimaging studies focused primarily on the importance of persistent activity in the PFC. Until recently, the potential importance of posterior cortical areas in WM had been largely neglected. Indeed, neural activity in monkey inferotemporal cortex is less robust during memory delays and the selectivity of activity appears to be disrupted by intervening distractors, while PFC representations appear resistant to distraction (Miller et al., 1996). Similarly, memoranda-specific delay period activity of neurons in the monkey PFC resists the effect of distractors, especially when compared to neurons in posterior parietal cortex (di Pellegrino and Wise, 1993; Constantinidis and Steinmetz, 1996; Suzuki and Gottlieb, 2013). Inferences stemming from the underlying logic of these distractor studies imply that the PFC, rather than posterior cortical areas, is critical for WM storage.

Recent human neuroimaging studies have further addressed this issue using various decoding methods, with a primary focus on whether information about remembered features can be found in visual cortex in the presence of an intervening distracting stimulus. Bettencourt & Xu (2016) decoded orientations held in visual WM using activation patterns in visual and parietal cortex on trials with and without distracting visual stimuli during the delay period (faces and gazebos). When it was predictable whether a distractor would or would not appear on a given trial, remembered orientations could not be decoded based on visual cortex activation patterns, but decoding from parietal cortex was successful. However, when distractor presence was unpredictable, both visual and parietal cortex represented remembered orientations during both distractor-present and -absent trials. Several subsequent commentaries (Ester et al., 2016; Gayet et al., 2018; Scimeca et al., 2018; Postle and Yu, 2020; Lorenc and Sreenivasan, 2021) and empirical reports (Lorenc et al., 2018; Rademaker et al., 2019; Hallenbeck et al., 2021) contested the theoretical importance of the null decoding performance in visual cortex for predictable distractors observed in Bettencourt & Xu (2016). The empirical studies largely replicated the finding in Bettencourt & Xu (2016) that activation patterns in parietal cortex contained information about WM content regardless of whether or not a distractor was present during the delay. However, decoded activation patterns in visual cortex do seem to depend on distractor presence, and alterations in these representations predict behavioral errors (Lorenc et al., 2018; Rademaker et al., 2019; Hallenbeck et al., 2021). Thus, persistent activity, as indexed by successful decoding of remembered information, survives visual distraction in many regions, and the impact of distraction on measured persistent activity in visual cortex is reflected in behavioral performance errors. These results across several studies suggest a critical role for stimulus-selective persistent activity in sensory cortex - it is often observed during delay periods, it appears unaffected by distractors when behavioral performance remains intact, and changes in persistent activity are reflected in changes in behavioral responses.

## Concluding remarks: persistent activity persists

Remarkably, for the past 50 years researchers studying the mechanisms of WM have used a variety of tools to characterize persistent activity across numerous types of memory tasks, across species, and across brain areas. It continues to stand as the central neural mechanism that supports WM. We have seen two arcs of research into persistent activity. One began in the front of the brain with single neuron recordings from the macaque PFC and led to what we have referred to as a canonical model of WM - a rich and mechanistically explicit computational model based in physiology and anatomy. The other began in the back of the brain using sophisticated machine learning algorithms that could precisely decode the contents of WM based on the patterns of neural activity in the human visual system. We’ve argued that such decoding is itself a manifestation of stimulus-selective persistent activity, just at a smaller scale than entire brain regions. Accordingly, persistent activity can be inferred not just from sustained elevated spiking of neurons, but from population level activity of fMRI BOLD signals sculpted by the content of memory.

The recent challenges to the canonical model include the various coding schemes (e.g., dynamic coding, mixed selectivity) and concerns about the evidence for persistent activity itself (e.g., artifacts, activity silent mechanisms). Another implicit challenge revolves around how “PFC-centric” the field has been when considering the neural mechanisms of WM. For instance, even if we accept each of these criticisms, the canonical model in its simplest form would need *no* revision if it was merely applied to brain areas other than the PFC. For instance, translating the classic findings of Funahashi - firing of neurons in macaque dorsolateral PFC persist over delays (Funahashi et al., 1989) and damage to this region impacts WM (Funahashi et al., 1993a) - to humans only requires shifting the locus from dorsolateral PFC to a brain region a bit more posterior in the precentral/arcuate sulcus (Mackey et al., 2016b). The various complexities with the types of coding and reliability of persistent activity in monkey PFC all disappear if the canonical model is instead applied to monkey areas like FEF and LIP (Hart and Huk, 2020). In those areas, single neurons clearly persist on single trials and the neurons form populations that represent WM features exactly as modeled (Wang, 2001) without the complexity of mixed selectivity and dynamic coding. Perhaps while we attempt to reconcile new discoveries about the PFC, we do not need to update our canonical model of WM. Overwhelmingly, the evidence indicates that simple, stable persistent activity among neurons in stimulus selective populations is the fundamental mechanism by which we maintain WM representations.

Moving forward, there are a number of questions that we instead need to address. Most relevant is: what, then, *is* the role of the dorsolateral PFC? Note that persistent activity is simply an observation and is not synonymous with WM maintenance (Curtis and Lee, 2010). Perhaps persistent activity in PFC reflects not the storage of WM features, but rather some mechanism related to the control of WM representations stored elsewhere, maybe by their own persistent activity (Miller and Cohen, 2001; Curtis and D’Esposito, 2003; Emrich et al., 2013; D’Esposito and Postle, 2015; Postle and Yu, 2020). Also, why do we see evidence of persistent activity, even for a simple single item WM task, in so many cortical and subcortical brain areas (Figs. 3–4; Christophel et al., 2017; Leavitt et al., 2017)? Redundancy is good to a point, but future research should try to figure out which of these numerous areas are necessary, what types of features they might be representing, and if they might be encoding different representational formats of WM. For example, disrupting persistent activity with intervening distraction (Lorenc et al., 2018; Rademaker et al., 2019; e.g., Hallenbeck et al., 2021) or TMS (e.g., Mackey and Curtis, 2017; Rademaker et al., 2017) may be able to disentangle the relative roles of different cortical regions. However, such efforts are tricky, as a distractor may not affect a top-down control signal, especially when passively-viewed. Experiments parametrically manipulating task demands in concert with visual distraction may help further clarify the relative role different brain regions play in WM tasks. An especially promising avenue for future exploration is comparing decoded feature values from single trials of fMRI activation to behavioral errors on those same trials (Li et al.; Ester et al., 2016; Hallenbeck et al., 2021).

WM is one of the few higher-level cognitive systems that we have made substantial progress towards understanding its neural implementation. Persistent activity has been at the heart of this success. While it is inevitable that additional mechanisms will be discovered, we have little doubt that persistent activity will persist as the primary explanation for how neural systems maintain WM representations

## Acknowledgments/Funding

CEC: R01 EY-016407 and R01 EY-027925 and TCS: F32-EY028438; Sloan Research Fellowship

